# CyChat: a conversational Cytoscape app for no-code, reproducible network analysis

**DOI:** 10.64898/2026.08.28.747833

**Authors:** Jeanine Liebold, Merle Stahl, Jan-Ole Schulze, Mohammad Mehdi Razavi, Gary D. Bader, Stefan Kurtz, Jan Baumbach

**Affiliations:** Institute for Computational Systems Biomedicine, Universität Hamburg, Hamburg, Germany; Faculty of Mathematics, Informatics and Natural Sciences, Center for Bioinformatics, Universität Hamburg, Hamburg, Germany; Donnelly Centre for Cellular and Biomolecular Research, University of Toronto, Toronto, ON, Canada; Department of Mathematics and Computer Science, University of Southern Denmark, Odense, Denmark

**Keywords:** Cytoscape, Large Language Models (LLMs), AI agents, Network analysis, Reproducibility

## Abstract

Network-based analyses of molecular interactions are useful for interpreting high-throughput omics data and identifying therapeutic targets. Cytoscape is the standard platform for these tasks, but users face a trade-off between accessible graphical workflows that are difficult to document and reproducible automation in Python or R that requires programming expertise. General-purpose coding assistants can generate Cytoscape Automation scripts, but remain external to Cytoscape.

We present CyChat, a Cytoscape Desktop app that integrates a chat interface and a large language model (LLM) agent into the application. CyChat translates natural language into executable Cytoscape Automation workflows, runs generated Python code, and exports chat sessions with executed code as standalone Jupyter notebooks. To reduce setup barriers, CyChat includes an embedded Python runtime and supports both cloud-based and locally hosted LLMs.

CyChat was evaluated across ten Cytoscape workflows using seven LLM providers, each represented by one LLM. The strongest configuration achieves a pass rate above 99%. In a qualitative evaluation based on a published network visualization, CyChat completes the task in 1.5–5 minutes, compared with 15–20 minutes for manual GUI workflows by computational biologists. CyChat is available through the Cytoscape App Store at https://apps.cytoscape.org/apps/cychat.

## 1 Introduction

Network analysis is central to interpreting high-throughput omics data and prioritizing therapeutic targets. Cytoscape [1] is the standard platform for this work and one of the most widely cited tools in network biology. Most users interact with Cytoscape through its graphical user interface (GUI), accessible via desktop and web browser [2]. While GUI-based workflows are accessible to domain experts, they are often difficult to reproduce because the interactions delivering the network were not captured [3].

Manual scripting through Cytoscape Automation [4] in R or Python enables reproducible workflows, but requires programming expertise that many domain experts lack. Recent tools improve selected network-analysis steps or data sources, but do not provide a general conversational interface for reproducible Cytoscape workflows.

AI coding assistants offer another route by generating Cytoscape Automation code, or can be connected to a live Cytoscape session through a Cytoscape Model Context Protocol server [5]. However, these solutions remain external to Cytoscape and are oriented toward users who are willing to switch between Cytoscape and a separate chat client or execute scripts in terminals and notebooks. This limits their usefulness for GUI-first users who prefer to perform analyses directly within Cytoscape. To our knowledge, no agent runs natively inside the Cytoscape Desktop application with full chat integration and automatic workflow execution.

We introduce CyChat, a Cytoscape Desktop app that embeds a large language model (LLM) agent directly inside the application. CyChat translates natural-language requests into executable Cytoscape Automation workflows and combines four technical features: (i) a provider abstraction for six cloud-based LLM providers and locally hosted LLMs through Ollama, (ii) a self-contained bundle with an embedded Python runtime for macOS, MS-Windows, and Linux, (iii) one-click export of chat sessions as reproducible Jupyter notebooks containing user messages, executed code, and CyChat responses, and (iv) a three-layer Python sandbox that restricts file access to read and write operations performed by py4cytoscape and pandas.

To assess CyChat’s effectiveness and efficiency, we built a quantitative benchmark of ten representative Cytoscape tasks, each with an automatic success check, and ran it across seven LLM providers. In the best configuration, CyChat passes over 99% of these runs.

## 2 Data and Methods

### 2.1 CyChat architecture

CyChat is implemented as a Cytoscape Desktop 3.9+ app (Java 17 OSGi bundle) with a self-contained Python runtime, requiring no separate interpreter or package manager. Its dockable chat panel connects to a backend that translates natural-language requests into Cytoscape Automation workflows and executes them in the live Cytoscape session via py4cytoscape and CyREST [6]. The architecture is shown in Figure 1.

**Figure 1:**
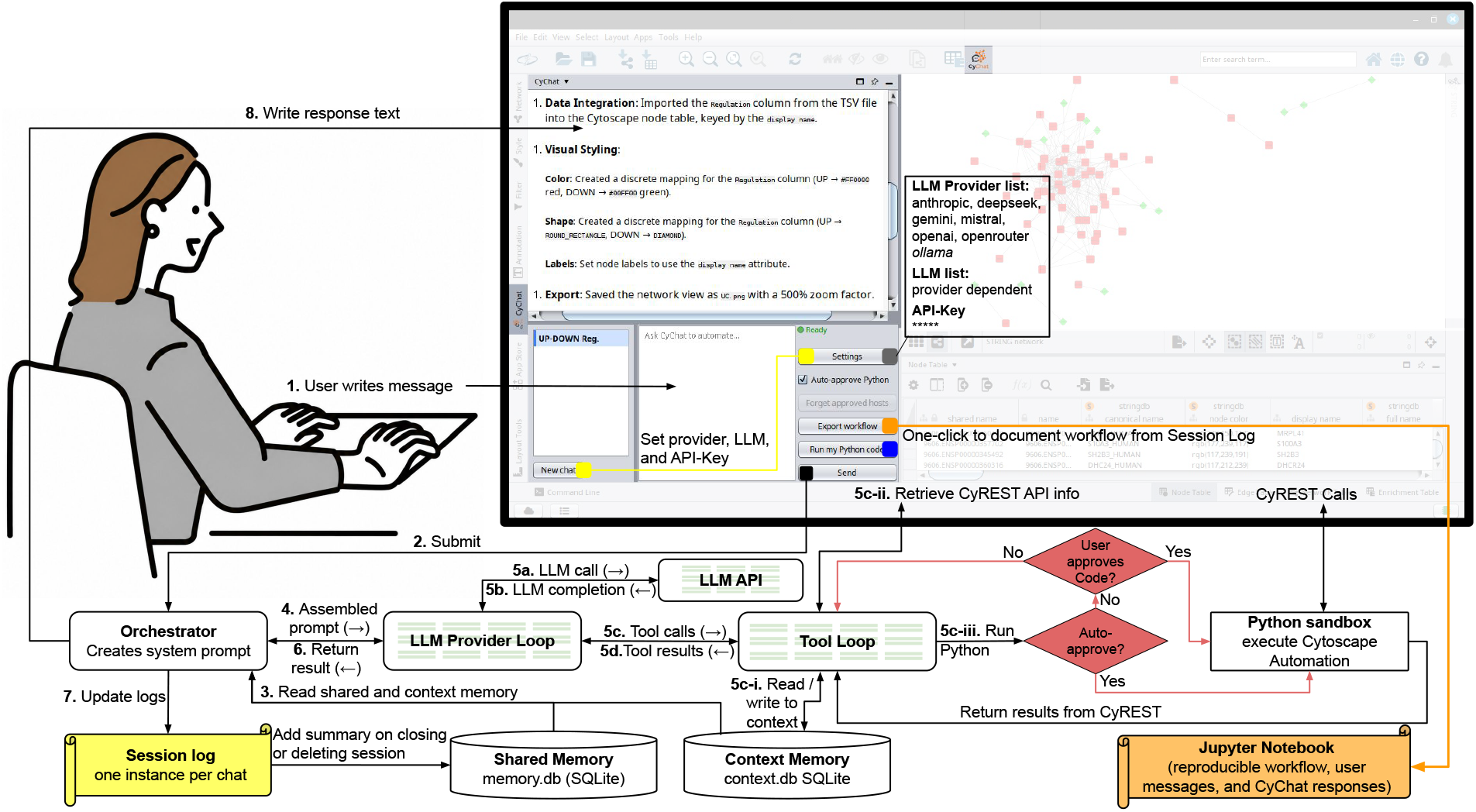
Architecture and workflow of CyChat. The figure shows the Cytoscape-embedded chat panel, the orchestrator, provider-specific LLM/tool loops, sandboxed Python execution, memory, session logging, and notebook export. Numbered arrows indicate the flow from user message to prompt assembly, LLM/tool execution, response generation, and session-log update. Colored UI controls start a new chat with user-defined settings (yellow), edit these settings (grey), export the workflow (orange), or run user-provided Python code (blue). Web requests require approval, and approved sites are remembered.

In the backend, CyChat combines a fixed system prompt with dynamic context that is refreshed for each user message. The fixed prompt defines the agent role, available tools, decision rules, execution protocol, coding constraints, and error-recovery strategy. An *Orchestrator* builds the assembled prompt from the user message, current session context, and shared memory. The prompt is then passed to the provider-specific agentic loop (Figure 1, shaded nodes). The session context summarizes the live Cytoscape state, recent errors, conversation history, and tool-created variables, while shared memory stores corrections, preferences, error patterns, and reusable workflow facts in a database. The assembled prompt is passed to an LLM provider-specific loop that adapts tool calls and responses to provider-specific request and response formats of six cloud-based LLM providers (Anthropic, OpenAI, Google Gemini, Mistral, DeepSeek, and OpenRouter) and Ollama. During this loop, CyChat sends the request to the selected LLM, executes generated tool calls when required, returns tool outputs to the LLM, and repeats this process until a final response is produced. CyChat is able to query the CyREST API for information about installed apps at runtime. Thus, automation coverage reflects the user’s installation rather than a hardcoded task set. In addition, the agent supports user guidance by answering GUI-navigation questions (e.g., “where is network analyzer?”) in text only, without generating or executing code.

### 2.2 Sandboxed execution and Cytoscape workflow documentation

The execution tool `run_python` passes through an approval gate. When auto-approval is disabled, each proposed snippet is shown as an editable chat bubble that users can edit, decline, or approve before execution. Approved code is forwarded to a three-layer Python sandbox. First, an abstract syntax tree (AST) allow-list specifies which Python constructs, imports, and selected packages are permitted. Second, the restricted execution environment exposes only curated Python `_builtins_` and withholds primitives such as `eval, exec, open,` and `_import_`. Third, the bundled Python runner is extracted to a local cache and checked against the shipped binary before use on Linux, MS-Windows, and macOS. Together, these layers provide a defensive restriction mechanism against general file-system, network, subprocess, and dynamic-code operations while allowing approved py4cytoscape-based analysis.

Every user message and approved Python execution is recorded in the session log. The *Export workflow* button converts this log into a standalone Jupyter notebook containing user messages, executed code, and CyChat responses. The *Run my Python code* button lets users rerun or extend workflow steps in the same sandbox.

### 2.3 Benchmark design and evaluation

#### Quantitative benchmark

Ten representative Cytoscape tasks were selected from the Cytoscape Automation tutorial corpus (https://github.com/cytoscape/cytoscape-automation/wiki). The tasks cover data import, network construction, continuous mapping, subnetwork extraction, image and session export, network analysis, app installation, annotation, and column filtering. For each task, we wrote three wording variants, all sharing a common goal but with different phrasings of the user message. These variants were *Explicit* for complete, polite sentences, *Imperative* for direct commands, and *Fragmented* for broad, shorthand requests. Each combination was repeated five times. Together, seven LLM providers, ten tasks, three wording variants, and five replicates yielded 7 *×*10 *×*3 *×*5 = 1 050 tests (see Figure 2).

**Figure 2:**
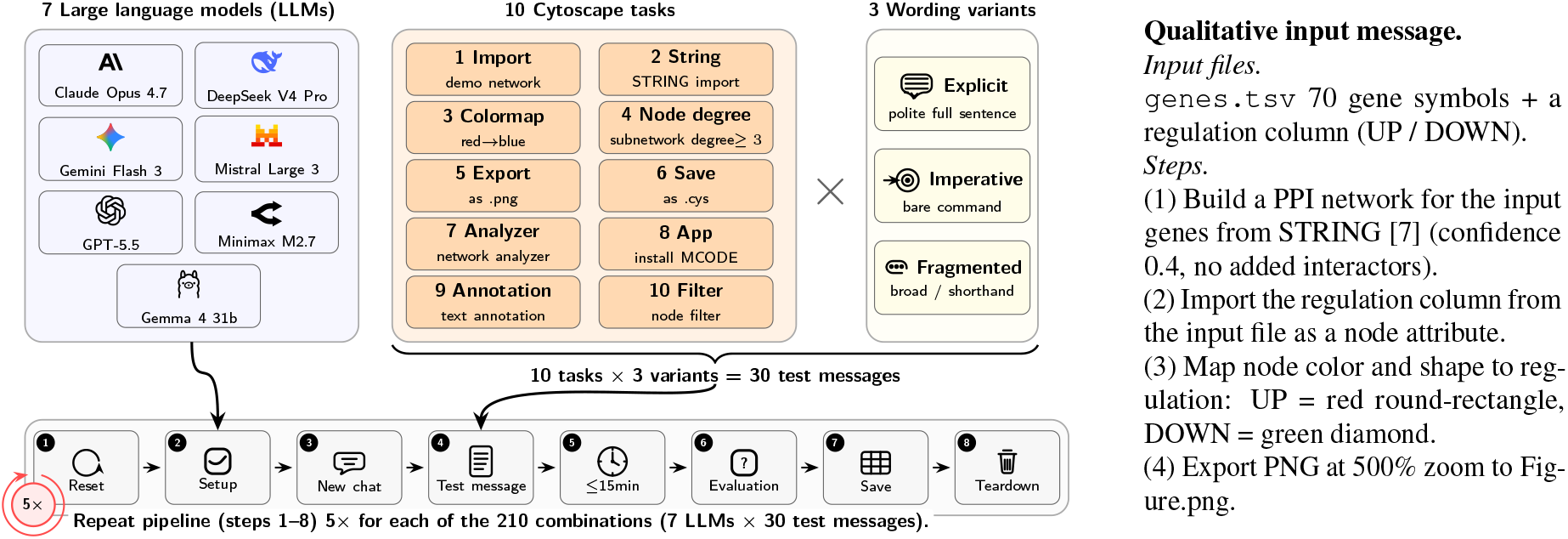
Quantitative and qualitative evaluation setup. Left panel: quantitative benchmark across LLMs (blue), Cytoscape tasks (orange), and wording variants (yellow), with five replicates per combination. Each run deletes CyChat’s shared memory and reopens the task session file (1), configures the selected LLM backend (2), opens a new chat (3), sends the task-specific message (4), waits up to 15 min (5), evaluates the result with py4cytoscape checks (6), saves logs (7), and tears down the chat (8). Right panel: the message used for the qualitative evaluation.

For each test, a message was submitted and the agent’s internal tool-call loop ran until termination or a 15 min wall-clock cap. The complete benchmark pipeline is shown in Figure 2. Pre-conditions were enforced by calling `close_session` followed by `open_session` on the task’s starting Cytoscape session (`.cys`) file. To ensure independent trials, both the session context and shared memory were cleared before every run, preventing any replicate from benefiting from the context or learned patterns of previous tasks. Remaining variation between replicates reflects provider-side inference behavior, including response sampling and any provider-side caching outside of our control. Post-conditions were evaluated programmatically using the Python package py4cytoscape against task-specific criteria recorded once when setting up the benchmark. Typical checks included file existence (e.g., an exported PNG or `.cys` file), comparison of node or edge counts against an expected count, and the color of nodes associated with extreme values (e.g., the lowest- and highest-`gal1RGexp` nodes for the continuous-mapping task). Each run yielded a binary pass/fail outcome, a wall-clock time, and an LLM API price. From these values, the three summary columns in Table 1 were computed. The *Reliable* column reports the number of tasks that an LLM passed reliably, defined as success in at least 12 of 15 runs, corresponding to 3 wording variants and 5 replicates. The time per passed test is the mean wall-clock duration over passing runs only, and the price per passed test is the mean API price over passing runs only. All runs were executed sequentially on a Linux Mint 22.3 laptop with an Intel Core i5-1145G7 CPU and a Cytoscape 3.10.4 instance.

**Table 1:**
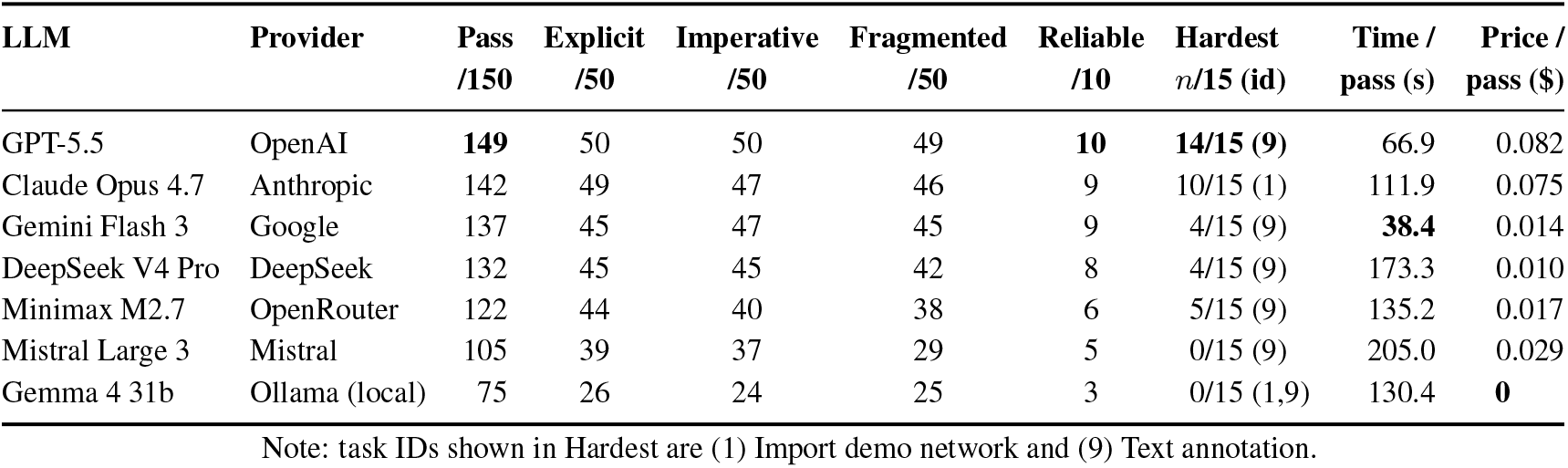
Quantitative benchmark results. **Pass** counts successful runs out of 150. **Wording columns (Explicit, Imperative, Fragmented)** count successful runs out of 50. **Reliable** counts tasks passed in at least 12*/*15 runs. **Hardest** reports the lowest per-task score and task ID. **Time/pass** and **Price/pass** are means over passing runs.

| LLM | Provider | Pass<br>/150 | Explicit<br>/50 | Imperative<br>/50 | Fragmented<br>/50 | Reliable<br>/10 | Hardest<br>n/15 (id) | Time /<br>pass (s) | Price /<br>pass (\$) |
| --- | --- | --- | --- | --- | --- | --- | --- | --- | --- |
| GPT-5.5 | OpenAI | **149** | 50 | 50 | 49 | **10** | **14/15 (9)** | 66.9 | 0.082 |
| Claude Opus 4.7 | Anthropic | 142 | 49 | 47 | 46 | 9 | 10/15 (1) | 111.9 | 0.075 |
| Gemini Flash 3 | Google | 137 | 45 | 47 | 45 | 9 | 4/15 (9) | **38.4** | 0.014 |
| DeepSeek V4 Pro | DeepSeek | 132 | 45 | 45 | 42 | 8 | 4/15 (9) | 173.3 | 0.010 |
| Minimax M2.7 | OpenRouter | 122 | 44 | 40 | 38 | 6 | 5/15 (9) | 135.2 | 0.017 |
| Mistral Large 3 | Mistral | 105 | 39 | 37 | 29 | 5 | 0/15 (9) | 205.0 | 0.029 |
| Gemma 4 31b | Ollama (local) | 75 | 26 | 24 | 25 | 3 | 0/15 (1,9) | 130.4 | **0** |

Note: task IDs shown in Hardest are (1) Import demo network and (9) Text annotation.

#### Qualitative evaluation

The Cytoscape visualization from Figure 4B of Zhang et al. [8] was chosen as an example workflow. This figure was published on February 13, 2026, which is after the training cut-offs of the two evaluated LLMs (Jan. 2025 for Gemini Flash 3 and Dec. 2025 for GPT-5.5). It is therefore unlikely that the figure could be reproduced from memorized training data. We extracted 70 genes and their up- and down-regulation labels from Zhang et al. [8]. Using this list, we described the task in natural language (cf. right panel of Figure 2).

Five figures based on this task description were created. Three computational biologists with experience in PPI network analysis (CB_1_–CB_3_, two novices and one occasional Cytoscape user) created the figure manually using the GUI. They received no assistance but were allowed to consult the Cytoscape documentation while performing the task. For all three, time was measured from the first Cytoscape interaction to the point when the figure was saved.

The task description was entered into CyChat, generating two figures: one using the Gemini Flash 3 backend and the other using GPT-5.5. The time from submitting the description to the final response was measured. The task was considered solved if the figure displayed a PPI network with 70 nodes and 305 edges, using rounded red rectangles for up-regulated genes (56 nodes) and green diamonds for down-regulated genes (14 nodes).

## 3 Results

### Quantitative benchmark

Table 1 shows the result of benchmarking CyChat across seven LLMs, ten Cytoscape tasks, three wording variants, and five replicates, yielding 1 050 runs. The strongest LLM, GPT-5.5, passes 149*/*150 runs, while four LLMs pass at least 130*/*150 runs.

Message wording has limited effect for most high-performing configurations. For five configurations, GPT-5.5, Claude Opus 4.7, Gemini Flash 3, DeepSeek V4 Pro, and Gemma 4 31b, the three wording-variant subtotals, Explicit, Imperative, and Fragmented, each out of 50, differ by at most three runs. The two exceptions, Mistral Large 3 (39*/*37*/*29) and Minimax M2.7 (44*/*40*/*38), perform worst on the Fragmented variant. Failures concentrate in specific tasks rather than across the full benchmark. Six of the seven configurations record their lowest task score on task 9 (text annotation). Gemma 4 31b ties at 0*/*15 on task 1 (import demo network) and task 9, while Claude Opus 4.7 records its lowest score on task 1 at 10*/*15. Runtime and prices differ substantially across configurations. Among passing runs, Gemini Flash 3 is fastest with a wall-clock time of 38.4 s, whereas Mistral Large 3 is slowest at 205.0 s. Mean API price per passing run is lowest for DeepSeek V4 Pro at $0.0095 and highest for GPT-5.5 at $0.0822.

### Qualitative results

All evaluated figures fulfilled the predefined success criteria. CyChat completed the task faster than CB_1_–CB_3_ (see Figure 3 for detailed completion times). The remaining differences were limited to visual appearance, i.e. color shades, node proportions, and layouts. These differences, however, occurred in outputs generated manually or by CyChat. They did not affect the encoded network structure or the interpretation of the regulation groups, and therefore reflect permissible Cytoscape styling choices rather than task failures.

**Figure 3:**
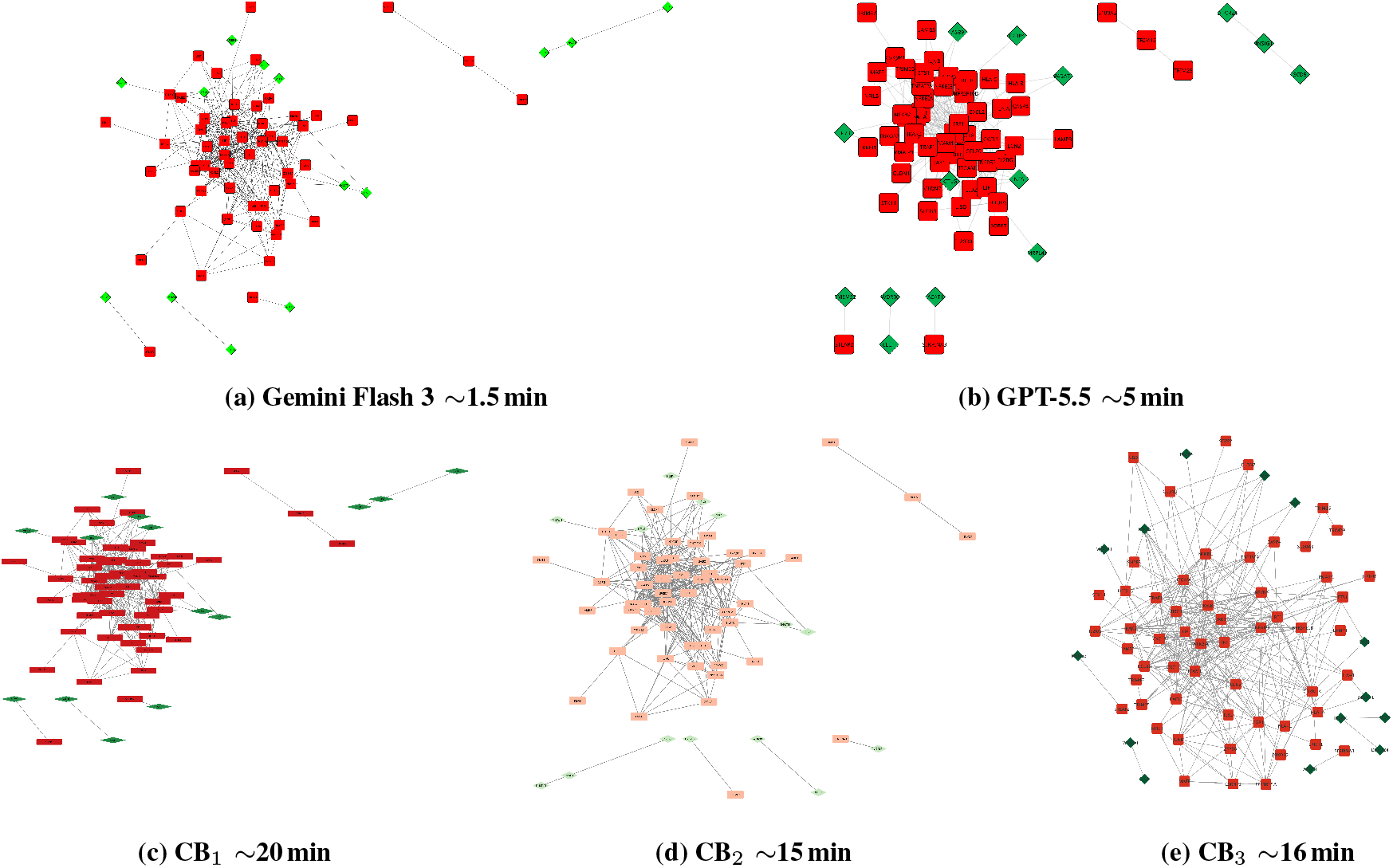
Qualitative evaluation based on Figure 4B in [8]. Per-panel labels report the LLM backend for CyChat in (a) and (b), the computational biologist ID in (c)–(e), and wall-clock time in all panels. All figures satisfy the predefined node, edge, color, and shape criteria (see Section 2.3, Qualitative evaluation).

## 4 Conclusion

CyChat narrows the gap between accessible GUI-based exploration and reproducible programmatic automation. Unlike external coding assistants, it embeds an LLM agent inside Cytoscape Desktop, letting GUI-first users express analysis goals in natural language, run them through Cytoscape Automation, and export approved executions as reusable Jupyter notebooks.

Our benchmark shows that this integration is technically feasible across diverse LLM providers and Cytoscape tasks. The strongest configuration passes 149*/*150 quantitative tests, and four configurations pass at least 130*/*150 tests. The qualitative evaluation shows that CyChat reproduced a recently published network visualization faster than the three manual GUI users tested while preserving the executed steps as code. CyChat is not intended to replace expert judgment. Its main advantages are speed and reproducibility, since every analysis is captured as executable code. This makes it easier to publish reproducible analyses, and allows new users to turn the informal description of a published workflow into code to reproduce it.

## Conflict of interests

No competing interests are declared.

## Funding

This work was supported by a fellowship within the IFI programme of the German Academic Exchange Service (DAAD).

## Notes

### Competing Interest Statement

The authors have declared no competing interest.

